# Cigarette smoke impairs pulmonary vascular function through nAChR activation

**DOI:** 10.1101/2024.05.20.594977

**Authors:** O Munar-Rubert, R Andreu-Martínez, J Rodríguez-Pérez, N López, B Barreira, E Fernández-Malavé, G Peces-Barba, C Muñoz-Calleja, A Cogolludo, MJ Calzada

**Affiliations:** Departament of Medicine, School of Medicine, Autónoma University of Madrid. Research Institute La Princesa Hospital (IIS-Princesa), Madrid, Spain; Departament of Pharmacology and Toxicology, School of Medicine, Universidad Complutense of Madrid, Madrid, Spain; Centro de Investigación Biomédica en Red de Enfermedades Respiratorias (CIBERES), Instituto de Salud Carlos III, Madrid, Spain; Department of Immunology, Ophtalmology and ENT, Complutense University of Madrid, Madrid, Spain; Respiratory Research Unit, Biomedical Research Unit, Health Research Institute Fundación Jiménez Díaz, Madrid, Spain

**Author notes:** These authors contributed equally to this work (O Munar-Rubert, R Andreu-Martínez).

## Abstract

Tobacco smoke is the main risk factor for the development of chronic obstructive pulmonary disease (COPD), a major health concern worldwide. Despite current therapies alleviate symptoms; there remain some limitations in the efficacy of treatments to curb COPD and its cardiovascular morbidities, particularly pulmonary hypertension. Our previous studies demonstrate that cigarette smoke (CS) has direct effects on pulmonary vascular tone homeostasis and contribute to pulmonary arterial dysfunction. This is in part due to altered activity of the voltage-dependent K^+^ channel, and to an exacerbated oxidative stress promoting a switch in the sGCs redox state. However, further characterization of the molecular basis of CS-mediated PA dysfunction is needed for more effective targeted treatment and prevention. Our current studies explored these molecular pathways and specifically addressed their contribution to the cellular contractile apparatus within pulmonary arteries. Our results proved deleterious effects on the contractile machinery of pulmonary artery smooth muscle cells. Increased oxidative stress and calcium dysregulation resulting from the activation of acetylcholine receptors (nAChR) in the pulmonary artery led to the manifestation of these effects. This groundbreaking discovery unveiled, for the first time, the expression of these receptors in human pulmonary arteries. Furthermore, we proved that inhibitors directed at these receptors demonstrate efficacy in alleviating various harmful effects of smoking and safeguarding pulmonary artery function from damage. These discoveries hold significant clinical implications, as they suggest that treatment with nAChR-targeted inhibitors could constitute a viable therapeutic option for COPD-related pulmonary hypertension in patients who do not respond to conventional medication.

## INTRODUCTION

Chronic obstructive pulmonary disease (COPD) is a complex respiratory pathology characterized by chronic bronchitis, emphysema, small airway obstruction, and persistent respiratory symptoms. Environmental factors, such as the rising prevalence of fine particulate matter in the environment and cigarette smoking (CS), exert a substantial influence on the development and progression of COPD (1). Some patients with COPD also experience alterations in resistance vessel structure mainly due to pulmonary artery (PA) remodeling, which in turn contributes to elevated pulmonary artery pressure (PAP) and the development of pulmonary hypertension (PH). This, in turn, leads to right ventricular dysfunction and, in many cases, to the death of the patient (2).

Despite its profound impact on public health, there remains a deficiency in curative and preventative therapies for COPD-related PH. Therefore, it is imperative to acquire a comprehensive understanding of the mechanisms underpinning this condition. While the direct effects of CS on the airways are well-documented, its impact on the pulmonary vasculature remains incompletely characterized. Conventionally, vascular remodeling was believed to result from alveolar hypoxia attributed to emphysema and airway obstruction. However, emerging evidence challenges this view, as elevated PAP only moderately correlates with hypoxemia and airflow limitation, and there is no correlation between elevated PAP and emphysema severity (3, 4). Additionally, previous studies show that mild COPD patients and smokers without airway obstruction exhibit media thickening in their pulmonary arteries, suggesting the involvement of other unknown mechanisms, presumably triggered by CS, in the development of PH (5). Our previous studies demonstrate that cigarette smoke extract (CSE) directly impairs PA vasoconstriction and vasodilation responses, indicating alterations intrinsic to the smooth muscle cell (SMC) layer. This was due in part to CSE-mediated dysregulation in the levels and activity of the voltage-dependent K^+^ channel Kv7.4 in pulmonary arteries (6).

Oxidative stress, understood as accumulation of ROS, can be caused not only by oxidizing substances such as tobacco, but also by an excessive production of these species or by a decrease in antioxidant defenses of the cells. This condition has been closely associated with COPD pathogenesis, as evidenced by increased oxidative stress markers in exhaled breath condensates and lung tissues of COPD patients compared to control subjects (7, 8). Similarly, alterations of the redox balance, observed for example in a decrease in glutathione availability (9), as well as dysregulated antioxidant response (10) explains why treatment with NAC, a glutathione precursor, is a widespread therapy for COPD patients (11, 12, 13). The excessive ROS production triggers a redox imbalance that contributes to organelle damage and disrupted homeostasis (14), as we have previously shown to occur with the NO-sGC signaling pathway in CSE-challenged human pulmonary artery smooth muscle cells (hPASMC) (10). These findings suggest that CS-induced changes in the PASMC may contribute to the development of PH in susceptible individuals. Furthermore, we have shown that CS induces mitochondrial dysfunction linked to a pulmonary artery vasodilation impairment (10). However, despite pulmonary artery contractile impairment has also been proved, its molecular mechanisms remain unclear and its relationship with oxidative stress should be considered.

Given the significant role of Ca^2+^ in contractile responses, it is plausible that CS may also affect the molecular components implicated in this signaling pathway. In this respect, chronic exposure of human airway cells and rat PASMC to CS induces the expression of plasma membrane Ca^2+^ channels (15, 16, 17). Several studies suggest that both nicotine (18) and tobacco smoke exposure (19), and in particular increased ROS (20, 21) have important effects on cytoskeleton structure and dynamics. Furthermore, changes in intracellular calcium signaling have also been involved in CSE-induced actin disorganization and dynamics (22). On the other hand, mitochondria also has an important role as a calcium-buffering organelle, and its dysfunction is often associated with calcium overload (23). Collectively, these results suggest a possible contribution of CS in decreasing contractile force generation in PA. Therefore, further characterization of the molecular basis of CS-mediated PA dysfunction is needed.

Smoking and nAChR (nicotinic acetylcholine receptors) share a close relationship, given that nicotine acts as an agonist for these channels. Recently, the expression of these channels has been demonstrated in the airway respiratory tract (24, 25). In this respect, some authors have highlighted the direct effects of nicotine on mitochondria. Although its mechanism of action remains unclear, data point to an involvement of nicotinic channels in these processes (26, 27). In line with this, previous reports suggest that calcium homeostasis may be affected during exposure to CS due to the influx of calcium and sodium ions through nAChR located in the plasmatic and the mitochondrial membrane (28, 26, 29). Because of these facts, it is plausible that signaling through these receptors would play an important role in the effects of smoking on the pulmonary artery.

This study aims to delve deeper into the molecular mechanisms underpinning the development of PH secondary to COPD. We hypothesize that CSE may be signaling in hPASMC and directly contributing to PA dysfunction in patients with COPD and COPD-related PH. Our results proved that tobacco exposure promoted oxidative stress, elevation of the intracellular calcium and alterations in the contractile machinery that explain CSE-induced alterations on PA contractility after CS exposure. These effects are promoted through nAChR signaling stimulated by nicotine present in the CSE. In this study, we conducted thorough research to investigate potential pharmaceutical applications aimed at alleviating symptoms associated with COPD-related pulmonary hypertension.

## MATERIALS AND METHODS

### Cell culture and treatments

Primary hPASMC were obtained from ScienCell Research Laboratories (#3110). The cells were grown in culture according to the specifications recommended by the manufacturer in SMC complete medium (ScienCell, #1101) containing 2% fetal bovine serum (ScienCell, #0010), 1X SMC growth serum (ScienCell, #2352 and #1152), 100 U/mL penicillin and 100 μg/mL streptomycin (ScienCell, #0503). Cells were grown to 90% maximum confluence, maintained at 37 °C in a humidified atmosphere of 5% CO2 and used for a maximum number of eight passages. In the indicated experiments, cells were pretreated with 25 nM of the mitochondrial antioxidant mitoTEMPO (Sigma, SML0737) or with 15 mM n-acetylcysteine (NAC) for 2 hours (Sigma, A7250) prior to co-treatment with CSE. Nicotine (Sigma Aldrich, N3876) was used at 2,5 μM for 30 minutes. This concentration of nicotine was based on the estimated concentration of nicotine present in the sputum of chronic smokers (found to be between 0.5 and 5 μM (30)). As a positive control for nAChR activation, cells were treated with 10 μM of the agonist N-[(3R)-1-Azabiciclo[2.2.2]oct-3-yl]furo[2,3-c]pyridine-5-carboxamide (PHA-543613, from Sigma, PZ0135) for 30 min. Other treatments were as follows: before CSE treatment cells were pre-treated with 50 μM Verapamil (VP) for 4 hours, with 50 μM 2-Aminoethyl diphenylborate (2-APB) for 2 hours, or 5 μM cytochalasin B (cytB) (Merk, C6762) for 2 hours. Afterwards cells media was washed out and replaced with fresh media containing CSE at the indicated concentrations. Pre-treatment was done with 10 μM Mecamylamine hydrochloride (Me) (Sigma Aldrich, M9020) for 2 hours, and 100 nM alpha bungarotoxin (α-bt) (Alomone, #B-100) for 30 minutes, and then CSE treatment for the indicated time.

### CSE preparation

CSE was prepared from commercial Kentucky 3R4F cigarettes (University of Kentucky Lexington, KY, 9.4 mg tar and 0.73 mg nicotine per cigarette). The smoke from the complete consumption of five cigarettes was continuously bubbled into 150 mL of PBS using VACU-SAFE aspiration system (INTEGRA Biosciences AG) at a vacuum pump flow rate of 8 L/min. CSE solution was filtered through a 0.20 μm-pore system, immediately aliquoted and kept at −80 °C until used. To ensure a similar preparation amongst different batches, CSE concentration was measured spectrophotometrically at 320 nm wavelength. This solution was considered to be 100% CSE and was diluted to obtain the desired concentrations for each experiment. To avoid the exposure to volatile substances from the smoke, untreated and CSE-challenged cells were always kept in different incubators.

### PCR analysis

Cells were grown to 90% confluence in culture plates and total RNA was isolated using TRI Reagent (Molecular Research Center Inc., TR118). RNA (1 μg) was reverse transcribed to complementary DNA (cDNA) with MultiScribeTM Reverse Transcriptase (Applied Biosystems, 4308228) in a final volume of 20 μL. cDNA (1 μL) was amplified with 1 μM of the specific primer pairs included in **Error! Reference source not found.**.

Electrophoresis was performed through agarose gels (0.7% agarose) prepared in tris-acetate (40 mM)/EDTA (10 mM) buffer and 5 μl of GelStar Nucleic Acid Stain 10,000 x (Lonza, 50535). 20 μl per sample resulting from PCR were mixed with Blue/Orange loading buffer loading dye 6x (PROMEGA, G190A) and loaded onto gel. Electrophoresis was run at 120V and gel bands were photographed with a transilluminator (Uvitec, BTS-20.M).

### Western Blot analysis

Cells were grown to 90% confluence in culture plates in the presence or absence of CSE for 24 h and lysates were prepared in non-reducing 2X Laemmli buffer, boiled at 95◦C for 10 min in the presence of 20 mM DTT, electrophoretically separated by SDS-PAGE and transferred onto 0.45-μm nitrocellulose membranes (GE Healthcare Life Sciences, 10600003). Total protein bands were reversibly stained with Fast Green FCF (Sigma, F7252) and imaged on ImageQuant LAS 4000 or Amersham ImageQuant 800 (GE Healthcare Life Sciences) for total protein quantification. Transferred proteins were incubated overnight at 4◦C with specific primary antibodies (Table 2). Horseradish peroxidase conjugated secondary antibodies (table 2) were added for 1 h at room temperature and protein signal was then visualized using Immobilon Forte (Millipore, WBLUF0500) on ImageQuant LAS 4000 or Amersham ImageQuant 800. Specific protein bands intensity was quantified by densitometry using ImageJ 1.51 software (U. S. National Institutes of Health, Bethesda, Maryland, USA) and normalized to the intensity of Fast Green FCF staining from each complete gel lane.

**Table 1:**
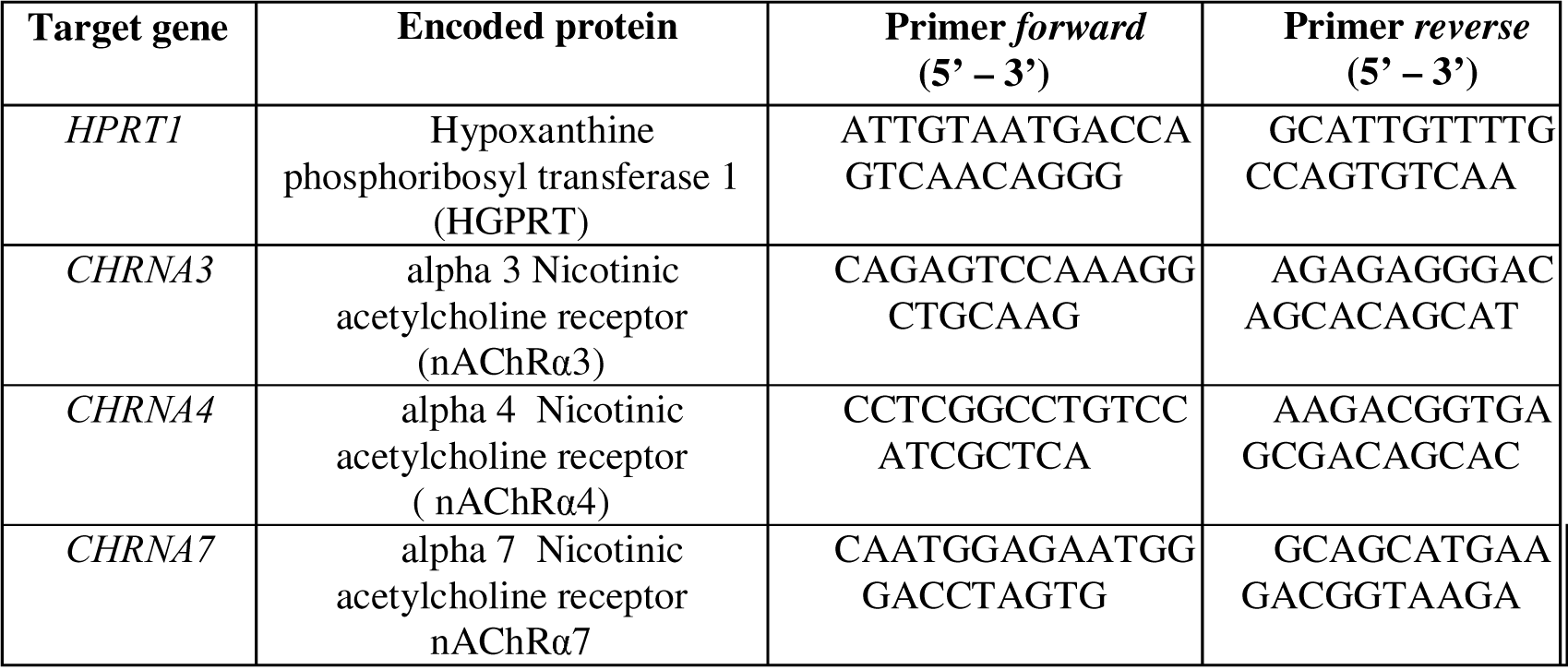
List of primer pairs used for gene expression analysis by PCR.

**Table 2:**
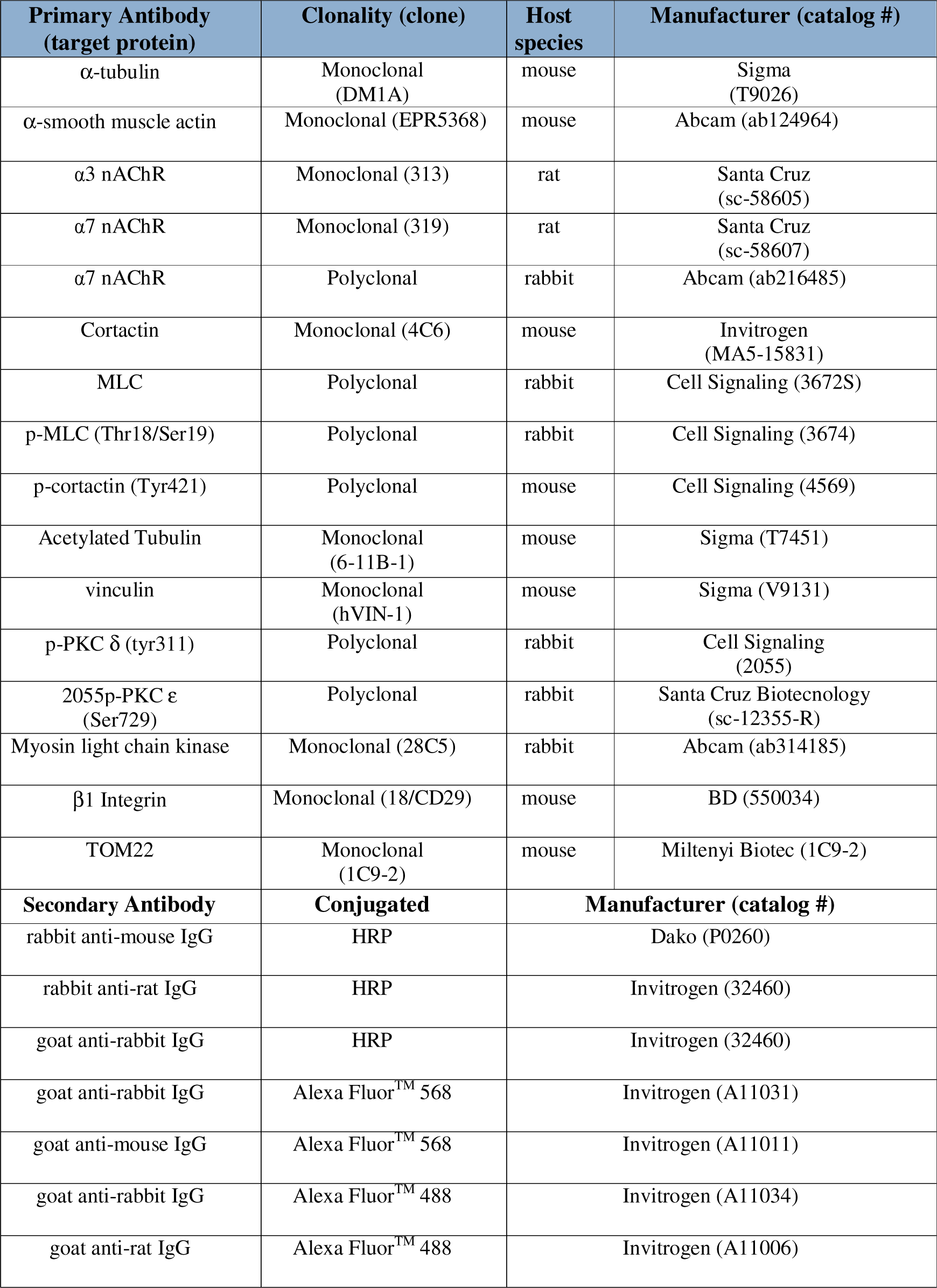
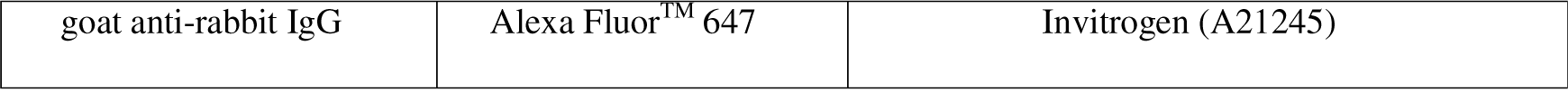
List of primary and secondary antibodies used for western blot and immunofluorescence, and their corresponding product information.

| Primary Antibody<br>(target protein) | Clonality (clone) | Host species | Manufacturer (catalog #) |
| --- | --- | --- | --- |
| α-tubulin | Monoclonal (DM1A) | mouse | Sigma (T9026) |
| α-smooth muscle actin | Monoclonal (EPR5368) | mouse | Abcam (ab124964) |
| $\alpha 3$ nAChR | Monoclonal (313) | rat | Santa Cruz (sc-58605) |
| $\alpha 7$ nAChR | Monoclonal (319) | rat | Santa Cruz (sc-58607) |
| $\alpha 7$ nAChR | Polyclonal | rabbit | Abcam (ab216485) |
| Cortactin | Monoclonal (4C6) | mouse | Invitrogen (MA5-15831) |
| MLC | Polyclonal | rabbit | Cell Signaling (3672S) |
| p-MLC (Thr18/Ser19) | Polyclonal | rabbit | Cell Signaling (3674) |
| p-cortactin (Tyr421) | Polyclonal | mouse | Cell Signaling (4569) |
| Acetylated Tubulin | Monoclonal (6-11B-1) | mouse | Sigma (T7451) |
| vinculin | Monoclonal (hVIN-1) | mouse | Sigma (V9131) |
| p-PKC $\delta$ (tyr311) | Polyclonal | rabbit | Cell Signaling (2055) |
| 2055p-PKC $\epsilon$ (Ser729) | Polyclonal | rabbit | Santa Cruz Biotechnology (sc-12355-R) |
| Myosin light chain kinase | Monoclonal (28C5) | rabbit | Abcam (ab314185) |
| $\beta 1$ Integrin | Monoclonal (18/CD29) | mouse | BD (550034) |
| TOM22 | Monoclonal (1C9-2) | mouse | Miltenyi Biotec (1C9-2) |
| Secondary Antibody | Conjugated | Manufacturer (catalog #) |  |
| rabbit anti-mouse IgG | HRP | Dako (P0260) |  |
| rabbit anti-rat IgG | HRP | Invitrogen (32460) |  |
| goat anti-rabbit IgG | HRP | Invitrogen (32460) |  |
| goat anti-rabbit IgG | Alexa Fluor<sup>TM</sup> 568 | Invitrogen (A11031) |  |
| goat anti-mouse IgG | Alexa Fluor<sup>TM</sup> 568 | Invitrogen (A11011) |  |
| goat anti-rabbit IgG | Alexa Fluor<sup>TM</sup> 488 | Invitrogen (A11034) |  |
| goat anti-rat IgG | Alexa Fluor<sup>TM</sup> 488 | Invitrogen (A11006) |  |

|  |  |  |
| --- | --- | --- |
| goat anti-rabbit IgG | Alexa Fluor™ 647 | Invitrogen (A21245) |

### Determination of total and mitochondrial O2^-^ and intracellular calcium

Total and mitochondrial superoxide levels, as well as intracellular calcium levels, were analyzed with fluorescent probes 10 μM dihydroethidium (DHE) (Invitrogen, D11347), 5 μM MitoSOX^TM^ Red (Thermo Fisher Scientific, M36008) and 1 μM Fluo-4 AM (ThermoFisher Scientific, F14201) respectively. Afterwards, cells were trypsinized and quantified by flow cytometry.

### Immunofluorescence

hPASMC were seeded onto fibronectin (20 μg/mL) coated 13-mm glass coverslips and then incubated under the desired experimental conditions. Cells were then fixed (20 minutes at 4°C) with 4% paraformaldehyde in PBS, and permeabilized (10 minutes RT) with 0,3% saponine in 0,2% BSA/PBS. Afterwards cells were labelled with fluorescence primary antibodies (Table 2) followed by Alexa Fluor^TM^ 488 and 568 labelled secondary antibodies and Phalloidin Alexa Fluor^TM^ 647 (Invitrogen, A22287) staining. Cells mounted in coverslips with Faramount Aqueous Medium were imaged with a Leica DMR microscope using an immersion 40x objective (HCX PL APO 100X/1.40–0.7 oil CS, Leica) and illuminated with a mercury lamp (ebq 100, Lightning & Electronics JENA). Ten images per sample were collected using Leica DCF360FX camera and LAS V4.1 software. The coherency coefficient was quantified on the same images using Orientation J, a software package comprising various plugins designed for ImageJ and Fiji. The Vector Field software assessed the vector modulus, typically indicative of coherence (Advances in Anatomy, Embryology and Cell Biology, vol. 219, Springer International Publishing, ch. 3, 2016).

Human pulmonary artery segments from surgical procedures performed at the Hospital Universitario La Princesa were used to analyzed the expression of α7 nAChR. Tissue was completely embedded into OCT compound prior to cryostat sectioning. Cut frozen sections at 5 µm were mounted on poly-L-lysine-coated histological slides and immunofluorescence staining was performed. Samples were fixed with 4% formaldehyde and permeabilized with 0,1% and 0,3% Tx-100 in PBS for 10 min each. After blocking with 10% FBS for 1h at RT, samples were incubated overnight at 4°C with primary α7 nAChR (ab216485) antibody and subsequently for 1h with secondary antibody Goat anti-rabbit 647 at 1:400. An anti-fade mounting media with DAPI (Life Science Inc., St. Petersburg, FL, USA) was used to fix the coverslip to a slide. The slides were examined using the THUNDER Imager 3D fluorescence microscope (Leica Microsystems) and processed with LASX software.

### Cytoskeleton recovery assay

The actin cytoskeleton was destabilized by incubation for 2h with cytB (Sigma-Aldrich, C6762-1MG) (5 μM) followed by a 1h recovery period, during which, cells were cultured without or with 15% CSE exposure. Conditions requiring pretreatment with any drug (NAC, α-bt and Me) were treated before adding cytB. Treatment was withdrawn during destabilization with cytB for reinstatement during the recovery period. Finally, the cells were fixed, labeled with Phalloidin Alexa Fluor^TM^ 568 (Invitrogen, A12380) and analyzed by flow cytometry. The mean fluorescence intensity (MFI) result of cells whose cytoskeleton had been destabilized with cytB was normalized with respect to cells that had received the same treatment but without cytB, in order to obtain the percentage of repolymerized F-actin.

### Mitochondrial extraction

Mitochondria were isolated with the mitochondrial Isolation kit from Miltenyi Biotec (130-094-532) according to the specifications recommended by the manufacturer. The eluent was analyzed by WB using antibodies against the proteins of interest (α3 and α7 nAChR). In parallel, the mitochondria suspension was analyzed by flow cytometry with an APC and FITC-A labeled secondary antibody.

### Cell senescence assay

Cell senescence was measured with the Invitrogen™ CellEvent™ Senescence Green Flow Cytometry Assay Kit (C10840). Cells were treated with or without 15% CSE and the specified pre-treatments (Me or α-bt) for 48 hours. After this time, cells were trypsinized and washed 2 times with 1X PBS. Then, fixed in 4% paraformaldehyde for 10 minutes at room temperature, and stained with the CellEvent™ Senescence Green Probe diluted 1/1000 in CellEvent™ Senescence Buffer for 2 hours at 37°C. Cells were washed in 1X PBS with 1% BSA, then resuspended in 1X PBS and analyzed by flow cytometry.

### Cell hypertrophy assessment

To determine the hypertrophic phenotype, hPASMCs were subjected to different dilutions of CSE for 24 hours and subsequently cell size was assessed. Median forward scattering of light from Ar 488 nm laser was quantified on a FACSCanto™ II cytometer (BD Biosciences), excluding cell debris and doublets from the analysis.

### Transfection of siRNA

Human α3-nAChR (sc-37055) and α7-nAChR siRNA (sc-42532) were purchased from Santa Cruz, USA. Control fluorescent scRNA (SIC005) was purchased from Sigma-Aldrich. Transfection of siRNA was performed according to the specifications recommended by the manufacturer using a commercial transfection reagent (sc-29528) from Santa Cruz, USA. Cells were then analyzed by WB or flow cytometry.

### Flow Cytometry

Table 2 specifies the primary and secondary antibodies used for this subsequent analysis. The gating strategy began by selecting cells based on forward scatter and side scatter parameters, and singlets by forward scatter area (FSC-A) versus forward scatter height (FSC-H). In all analyses, cells were stained with Ghost DyeTM 780 to mark cell viability. For mitochondrial analysis, the column eluent was first labeled with the specific primary antibody for 1 hour at 4°C and then with the appropriate fluorescent secondary antibody. Mitochondria were selected according to their size and complexity, excluding cellular debris, and positive for the mitochondrial marker TOM22. Median fluorescence was measured on a FACSCanto II cytometer (BD Biosciences). Specific changes in MFI were quantified by subtracting the fluorescence of the unlabeled treated control, and normalized to the average MFI of the different conditions.

### Vascular contractility measurement

PA from WT C57BL/6J mice were carefully dissected free of surrounding tissue, cut into rings (1.8-2 mm length) and pre-treated with 10 μM Mecamylamine (2 hours) or 100 nM alpha bungarotoxin (Alomone, #B-100) (30 minutes). Afterwards, rings were overnight-exposed to 15% CSE. Vessel segments were mounted on a wire myograph in the presence of Krebs physiological solution. Contractility was recorded by an isometric force transducer and a displacement device coupled with a digitalization and data acquisition system (PowerLab, Paris, France). Preparations were firstly stimulated by rising the K^+^ concentration of the buffer to 80 mM in exchange for Na^+^. Vessels were washed three times and allowed to recover before a new stimulation. Arteries were then stimulated with increasing concentrations of serotonin (5-HT, from 10^–8^ to 3·10^–5^ M, Sigma), endothelium-dependent vasodilator acetylcholine (ACh, from 10^–9^ to 10^–5^ M, Sigma) or increasing concentrations of the sGC stimulator riociguat (from 10^–9^ to 10^–5^ M).

All animal experiments and procedures were carried out in accordance with the guidelines of the European Community Council Directive (86/609/EEC) and the Spanish Royal Decree 53/2013 based on the European Union normative (2010/63/UE). All protocols were approved by the Committee for Research and Ethics of the Universidad Autonoma of Madrid (PROEX 322/14).

### Statistics

Data are presented as the mean ± SEM or as interquartile range as well as the p95 and p5, as specified. Two-tailed Student’s or one-sample t-tests were used to compare two groups and one-way or two-way ANOVA tests followed by Dunnet’s post-hoc test were used when comparing three or more groups, according with the conditions of normality and homoscedasticity. In case the assumptions of normality and homoscedasticity were not accomplished, Mann-Whitney’s test was used to compare two groups and non-parametrical Kruskal–Wallis test followed by Dunn’s post-hoc test was used to compare three or more groups. Statistical analyses were performed on GraphPad Prism 9.0 software (San Diego, CA, USA) or with R version 4.3.3.

## RESULTS

### hPASMC exposed to CSE show reduced levels of contractile machinery proteins

Disruption of cytoskeletal proteins can have profound effects on cellular function, including alterations in contractility and vascular tone. Considering our previous results on the relevance of ROS in vascular impairment after exposure to cigarette smoke (6, 10), understanding how ROS specifically affect the contractile response in the vasculature could shed light on potential therapeutic targets for mitigating smoke-induced vascular impairment. Thus, we evaluated the impact of CSE and ROS on specific cytoskeletal proteins. To this aim hPASMC were exposed to 10% or 15% CSE for 24 hours and afterwards, protein levels were analyzed by Western blot. Our results showed that cytoskeletal proteins such as tubulin and cortactin, as well as their post-translationally modified active forms, acetylated tubulin and phosphorylated cortactin, were significantly decreased after CSE treatment. Surprisingly, the levels of actin, which is a major component of the cytoskeleton and plays a central role in cell structure, remained unchanged (Figure 1). These results suggested that CSE may selectively impact certain components of the cytoskeleton. Additionally, we analyzed the impact of smoke exposure on proteins of focal adhesions, since these are dynamic protein complexes playing a crucial role in mechanotransduction. Similarly, the levels of vinculin were significantly decreased in CSE-treated cells (Figure 1). Furthermore, analyzing the levels of myosin light chain (MLC) and its phosphorylated active form (p-MLC) provides crucial insights into the contractile state. Our results demonstrated that total MLC levels remained unaltered, while p-MLC were significantly decreased in CSE-treated cells. Most importantly, the observation that the decrease in p-MLC in CSE-treated cells was prevented in the presence of the antioxidant N-acetylcysteine (NAC) highlights the significant involvement of oxidative stress in mediating the impact of CSE on pulmonary artery smooth muscle cell contractility (Figure 2A). Consistent with these findings, we noted reduced levels of the MLC kinase (MLCK) and the upstream kinase PKC ε and PKC δ in our CSE-treated cells (Figure 2B).

**Figure 1.**
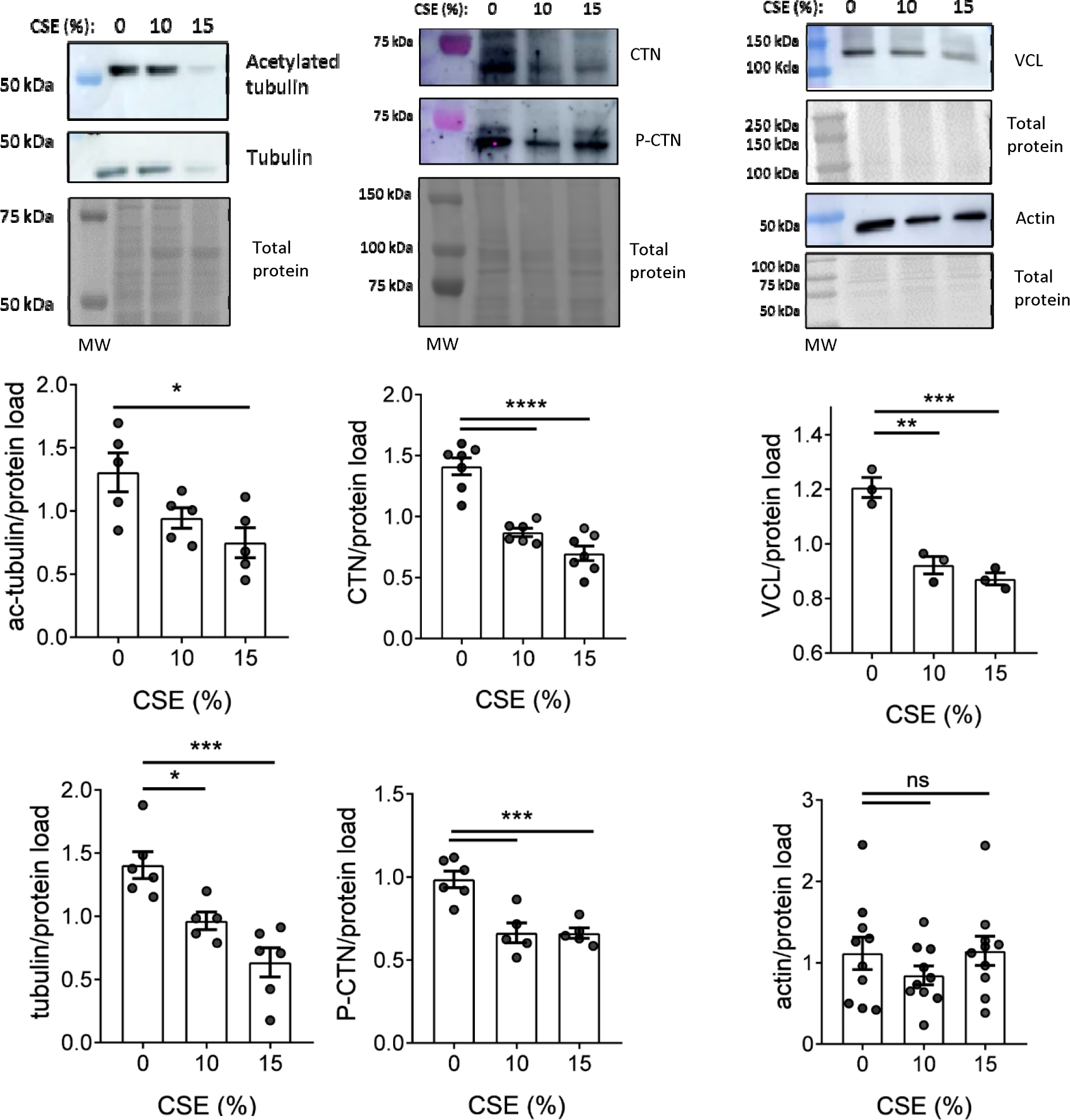
CSE effects on hPASMC contractile machinery proteins. hPASMCs were treated for 24 hours with 10% or 15% CSE as indicated and the levels of different and representative cytoskeleton proteins were analyzed by WB. From left to right and from top to bottom: acetylated tubulin, total tubulin, total cortactin (CTN), phosphorylated cortactin (p-CTN), vinculin (VCL) and total actin. For each protein analyzed, representative images showing the protein bands along with their respective quantifications by densitometry are presented. Densitometry measurements were adjusted to account for the intensity of total protein staining, serving as a loading control. Each data point corresponds to a single experiment, which is derived from the average of three technical replicates. Data are represented as the mean ± SEM; (n=3-10). Statistical analysis between groups was done by one-variable ANOVA test followed by Dunnet’s post hoc test (*)p<0.05, (**)p<0.01, (***)p<0.005, (****)p<0.0001, (ns) non-significant. MW: molecular weight marker.

**Figure 2.**
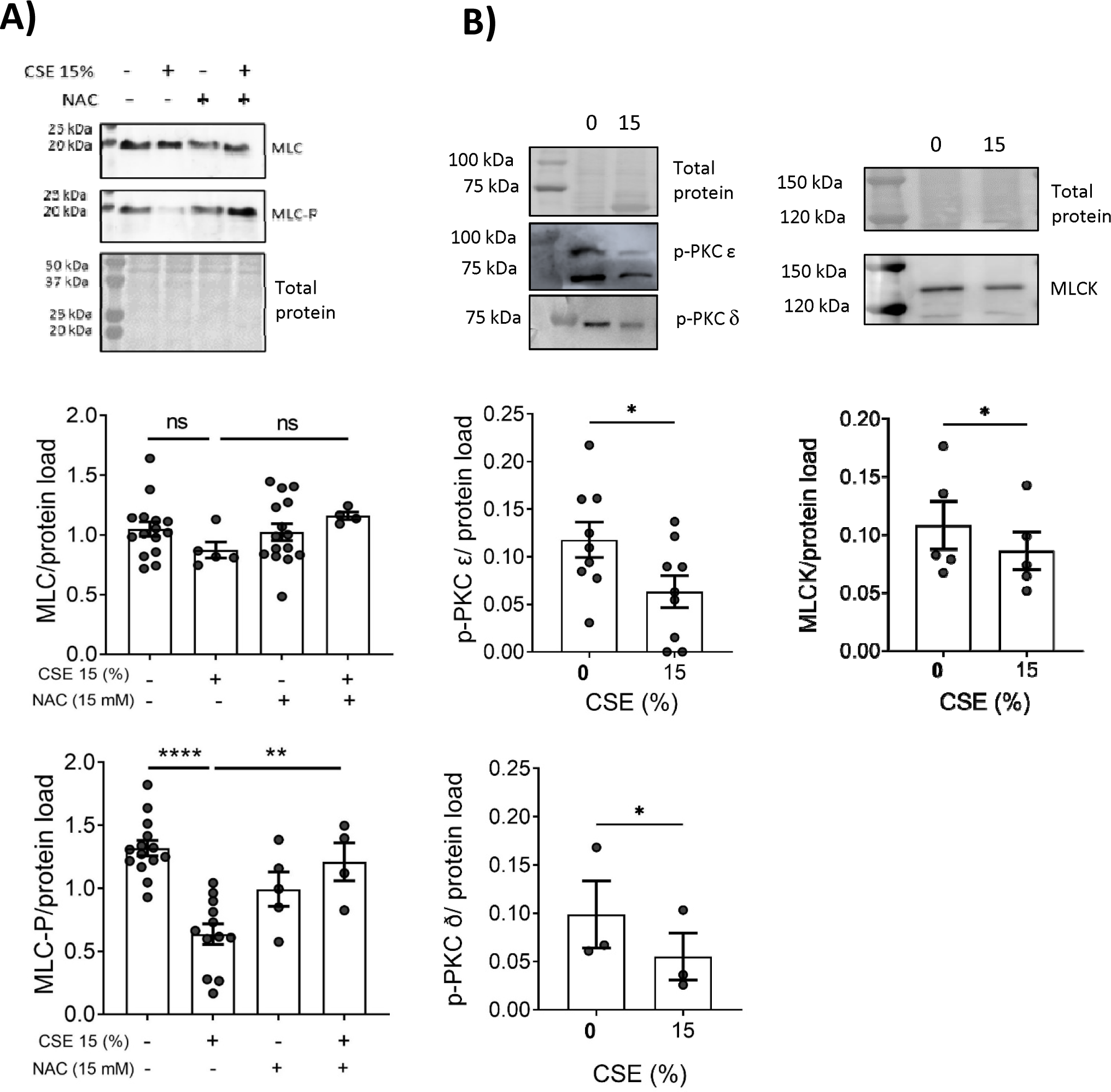
CSE effects on MLC phosphorylation and kinase levels. hPASMCs were treated for 24 hours with 0% or 15% CSE and when indicated cells were pretreated for 4 hours in the absence (-) or presence of 15 mM NAC (+) and maintained during CSE treatment. Afterwards the levels of the indicated proteins were analyzed by WB and quantified by densitometry **A)** Levels of myosin light chain (MLC) and its phosphorylated form (p-MLC) are shown. Data are represented as the mean ± SEM; (n=3-15). Statistical analysis between groups was done by one-way ANOVA test followed by Dunnet’s post hoc test (**)p<0.01, (****)p<0.0001, (ns)non-significant. MW: molecular weight marker. **B)** Levels of myosin light chain kinase (MLCK) and phosphorylated protein kinase C (p-PKCε and p-PKCδ isoforms). Densitometry measurements were adjusted to account for the intensity of total protein staining, serving as a loading control. Each data point corresponds to a single experiment, which is derived from the average of three technical replicates. Data are represented as the mean ± SEM; (n=3-9). Statistical comparisons between both groups were made using two-tailed Student’s t-test; (*)p<0.05. MW: molecular weight marker.

### CSE exposure promotes ROS-mediated functional alterations of the contractile system

Changes in actin filament morphology, density, or distribution patterns may indicate alterations in cytoskeletal dynamics and cellular function induced by CSE. To provide valuable information not just on the levels but also on the organization and distribution of actin filaments and focal adhesion within the cytosol we performed fluorescence microscopy staining with phalloidin, p-MLC and vinculin. Consistent with protein levels analyzed in Figure 1, the staining with phalloidin demonstrated actin bundles perfectly organized in control cells compared with CSE-treated cells, in which actin bundles showed more diffused and less organized. Furthermore, the analysis of the coherency coefficient, which indicates changes on bundles directions, demonstrated decreased coefficient in CSE-treated cells compared to control cells, this supporting a defect in cytoskeleton organization. Similarly, vinculin staining in CSE-treated cells showed alterations of focal adhesions characterized by a lower number of puncta arranged at the cell periphery, compared to control cells in which we observed a higher number of puncta evenly distributed in the cell cytoplasm (Figure 3A). This suggested a correlation between cytoskeletal remodeling and focal adhesion dynamics in hPASMCs.

**Figure 3.**
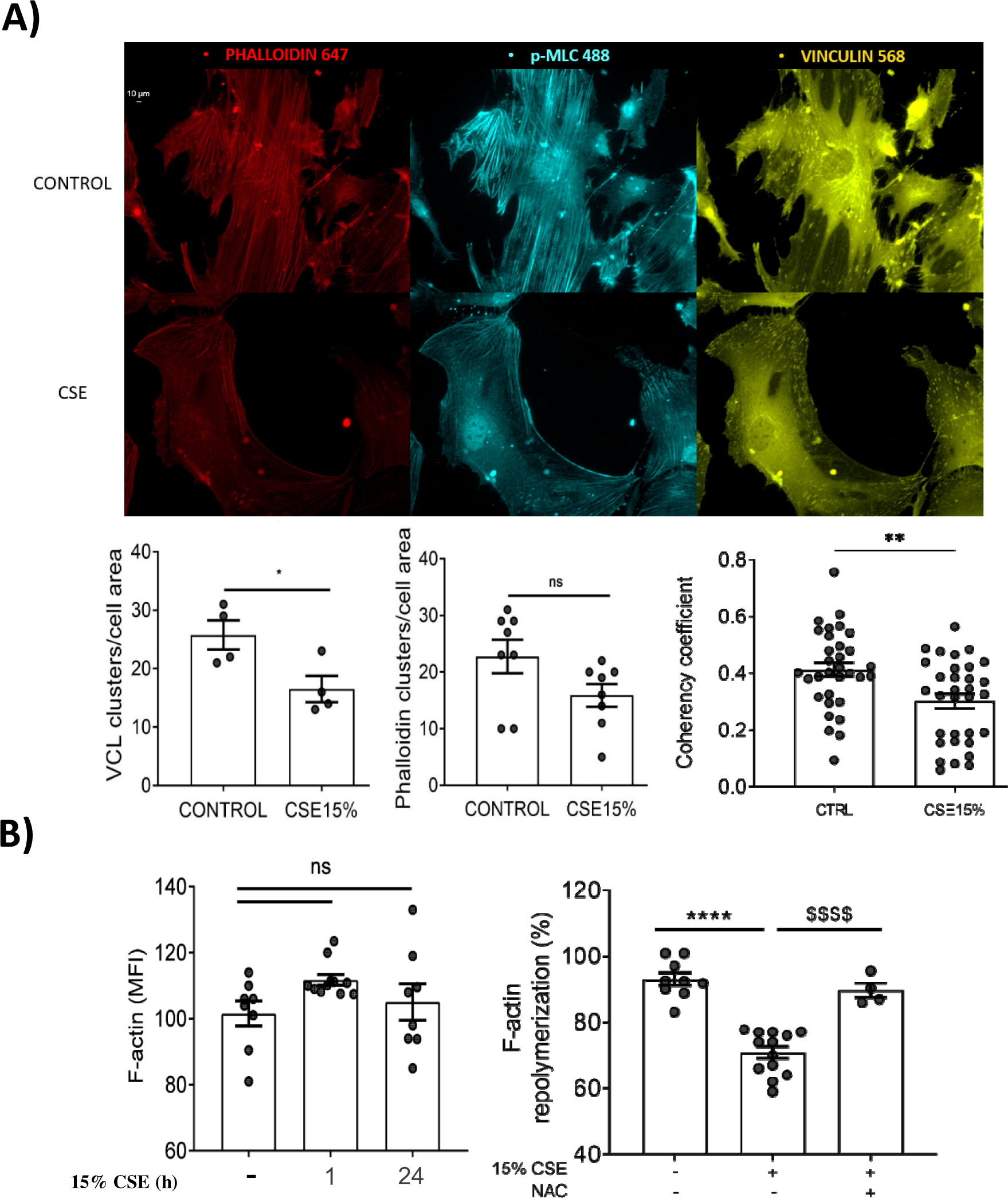
CSE effects on cytoskeleton organization and actin dynamics. Cytoskeleton organization and actin dynamics were examined by means of fluorescence microscopy and flow cytometry in hPASMCs following 1 or 24-hour treatment with 15% CSE **A)** Following cell treatment cytoskeletal organization was analyzed by fluorescence microscopy. Cell immunostaining with p-MLC (cyan) and vinculin (yellow) and phalloidin-Alexa FluorTM 647 (far red) staining as well as the quantification of phalloidin clusters/cell area and VCL clusters/cell area are shown. Scale bar = 10 μM. The coherency coefficient was also quantified on the same phalloidin stained images using Orientation J. At least 10 fields randomly selected and 4-5 cells from each condition were analyzed and representative images from a total of n=4-8 experiments performed are shown. A coherency coefficient close to 1 indicates a strongly coherent orientation of the local fibers in the direction of the ellipse long axis. A coherency coefficient close to zero denotes no preferential orientation of the fibers. **B)** Quantification by FACS of total F-actin levels (left) were analyzed in cells treated for 1 hour or 24 hours with 15% CSE staining with Phalloidin Alexa Fluor^TM^ 568. The percentage of F-actin repolymerization (right) was assessed in cells pretreated with or without 15 mM NAC for 4 hours. Afterwards cells were treated with 5 µM cytochalasin B for 2 hours and then allowed to recover for 1 hour with or without 15% CSE in the absence or presence of 15 mM NAC. The MFI of recovering cells was adjusted by the ratio between destabilized/non-destabilized cells. All data are represented by mean MFI ± SEM; (n= 3-13). Statistical comparison between groups was performed using one-way ANOVA test followed by Dunnet’s post hoc test. (ns) non-significant, (****)p<0.0001 points to significance between CSE treatment and control group, ($$$$)p<0.0001 points to significance within CSE group and NAC co-treatment.

In parallel actin dynamic experiments and quantification of F-actin levels were performed in cells treated for 1 hour with CSE after cytoskeleton destabilization with cytochalasin B and analyzed by flow cytometry. Interestingly, despite total F-actin levels were similar in control and CSE-treated cells (Figure 3B), actin polymerization recovery in cells treated with CSE was significantly decreased compared to untreated ones. Most importantly, the treatment with the antioxidant NAC prevented these effects (Figure 3B). These data suggest that the rise in reactive oxygen species generated following CSE treatment holds significant implications for the dynamics of the cell cytoskeleton, thereby contributing to the alteration of the contractile response.

### hPASMC express active **α**3 and **α**7 nAChR

Due to the observed alterations in the contractile machinery and the significance of calcium as a signaling molecule in vascular responses, we aimed to explore the contribution of calcium channels in this signaling process. Given that nicotinic receptors have not been documented in human pulmonary artery smooth muscle cells, we first analyzed their expression in these cells. Interestingly, this analysis unveiled, for the first time, the presence of both mRNA and protein levels of α3 and α7 channels in hPASMC (Figure 4A). Additionally, our results demonstrate for the first time the expression of these channels, in particular α7 nAChR in the three layers of human pulmonary artery samples (Figure 4B). Next, we evaluated CSE effects on increasing intracellular calcium and ROS levels. Our results showed that CSE treatment promoted a rapid and significant increase in intracellular Ca^2+^ levels, with its maximum peak at 30 min after treatment (data not shown). Moreover, this increase was parallel to the increase in total ROS levels (Figure 4C). It is worth mentioning that the activation of voltage-dependent Ca^2+^ channels is triggered by nicotine-mediated activation of nAChRs, this process facilitating Ca^2+^ influx. Thus, we explored if nicotine, as an important component of tobacco and an endogenous ligands for these channels, could replicate the effects observed with CSE. We observed that nicotine, as well as a commercial agonist for nAChR, PHA, induced an increase in Ca^2+^ and ROS levels that was comparable to that induced by CSE (Figure 4C). These findings provide further support for the hypothesis that CSE may exert its effects through the activation of nicotinic receptors in hPASMC.

**Figure 4.**
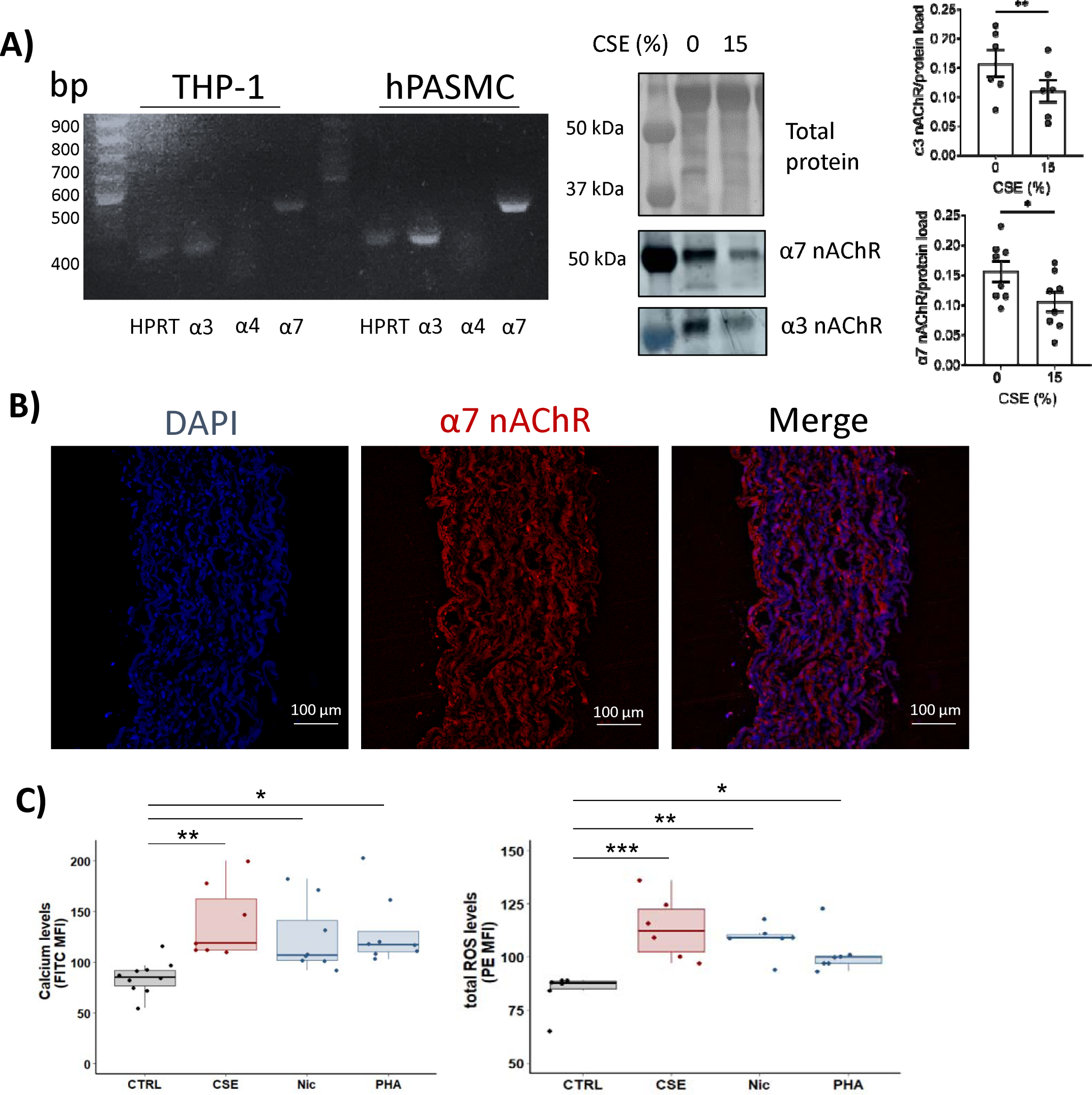
Analysis of nAChR levels and activity. **A)** Expression of nAChR subunits in human pulmonary artery smooth muscle cells or in THP-1 cell line (used as a positive control of the expression of these channels) was analyzed by PCR. Protein expression was quantified in presence or absence of 15% CSE for 24 hours. Densitometry measurements were adjusted to account for the intensity of total protein staining, serving as a loading control. Each data point corresponds to a single experiment, which is derived from the average of three technical replicates. Data represented as mean ± SEM; (n= 6-8). Statistical comparisons between both groups were made using two-tailed Student’s t-test; (*)p<0.05, (**)p<0.01. **B)** Immunofluorescence staining of α7 nAChR in human pulmonary artery. **C)** hPASMCs were treated for 30 min with 15% CSE or with nAChR agonists (2.5 µM nicotine or 10 µM PHA) as positive controls to confirm nAChR activity. Intracellular calcium and total ROS levels were measured with Fluo-4-AM and DHE probes respectively and analyzed by flow cytometry. Data are shown as interquartile range (p75 upper edge of box, p25 lower edge, p50 midline) as well as the p95 (line above box) and p5 (line below); (n=4-10). Statistical comparison between groups was performed using ANOVA test followed by Dunnet’s post hoc test; (*)p<0.05, (**)p<0.01, (***)p<0.005 points to significance between untreated and antagonist or CSE-treated group.

### Nicotinic receptor blockade prevents CSE-mediated increase in ROS and calcium

Our results have demonstrated that CSE exposure led to an increase in cytosolic ROS and calcium levels. To confirm whether blocking nAChR could prevent the rise in ROS and calcium induced by CSE, we analyzed the impact of alpha bungarotoxin (α-bt), a selective competitive antagonist for α7 nAChR, and mecamylamine (Me), a non-selective non-competitive nAChR antagonist, on calcium and ROS levels in hPASMC exposed to CSE. Our findings showed that treatment with either α-bt or Me significantly prevented the increase in cytoplasmic calcium and ROS levels caused by exposure to CSE, and these effects were noticeable as early as 30 minutes after treatment and lasted for at least 24 hours (Figure 5A). Next, we aimed to ascertain whether these effects were specific to α3 or α7 nAChR. To do so, we interfered their expression with specific siRNA for α3 or α7 nAChR and analyzed ROS and calcium levels. Treatment with the specific siRNAs resulted in partial interference, which was effective enough to induce a slight but significant reduction in calcium and ROS levels in CSE-treated hPASMCs (Figure 5B).

**Figure 5.**
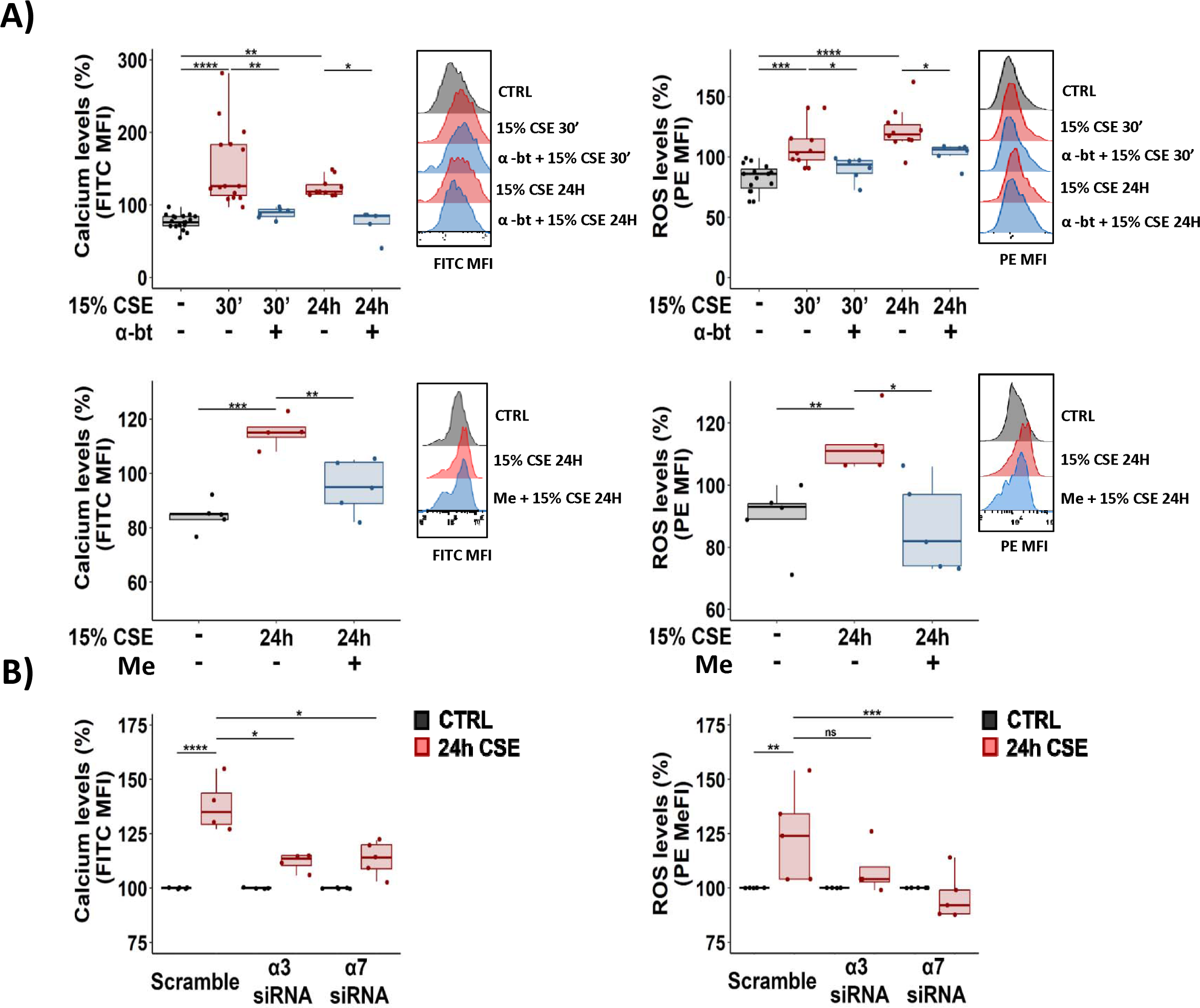
Effects of nAChR antagonists and interference on the elevation of total ROS and calcium levels induced by CSE. Intracellular calcium and total ROS levels were measured with Fluo-4-AM and DHE probes respectively and analyzed by flow cytometry. **A)** hPASMC were pretreated with nAChR antagonists: 100 nM α-bt (*top*) or 10 µM Me (*bottom*). Afterwards cells were cultured without (CTRL) or with 15% CSE at the indicated times. Data at the top are presented as interquartile range (p75 upper edge of box, p25 lower edge, p50 midline) as well as the p95 (line above box) and p5 (line below); (n=4-19). Representative half-overlay histograms of Fluo-4-AM and DHE fluorescence intensity are shown: control non-CSE treated (CTRL, *grey*), 15% CSE (*red*), α-bungarotoxin + 15% CSE or Me + 15% CSE (*blue*). **B)** Intracellular calcium and total ROS levels analysis in hPASMC interfered with specific siRNA for α3 or α7 nAChR and treated with 15% CSE for 24 hours. Data are presented as interquartile range (p75 upper edge of box, p25 lower edge, p50 midline) as well as the p95 (line above box) and p5 (line below); (n=4-5). Statistical comparison between groups was performed using one-way ANOVA test followed by Dunnet’s post hoc test; (*)p<0.05, (**)p<0.01, (***)p<0.005, (****)p<0.0001 points to significance between groups, (ns) non-significant.

### Isolated mitochondria from hPASMC express functional nAChR

As we observed previously, CSE led to an elevation in mitochondrial ROS levels, prompting us to assess whether the rise in mtROS induced by CSE could be attributed to the activation of nicotinic receptors. Our findings indicate that the increase in mtROS was effectively prevented solely by the hypertensive drug mecamylamine, not by α-bt. (Figure 6A). Considering that this toxin was not cell-permeant, this raised the possibility that nAChR were also present on the mitochondrial membrane of hPASMC. Consequently, we analyzed the expression of α3 and α7 nAChR in mitochondria. Our results revealed that both receptors were indeed expressed in mitochondria (Figure 6B). Therefore, their biological activity and connection to tobacco-induced mitochondrial dysfunction was confirmed after blocking them with mecamylamine.

**Figure 6.**
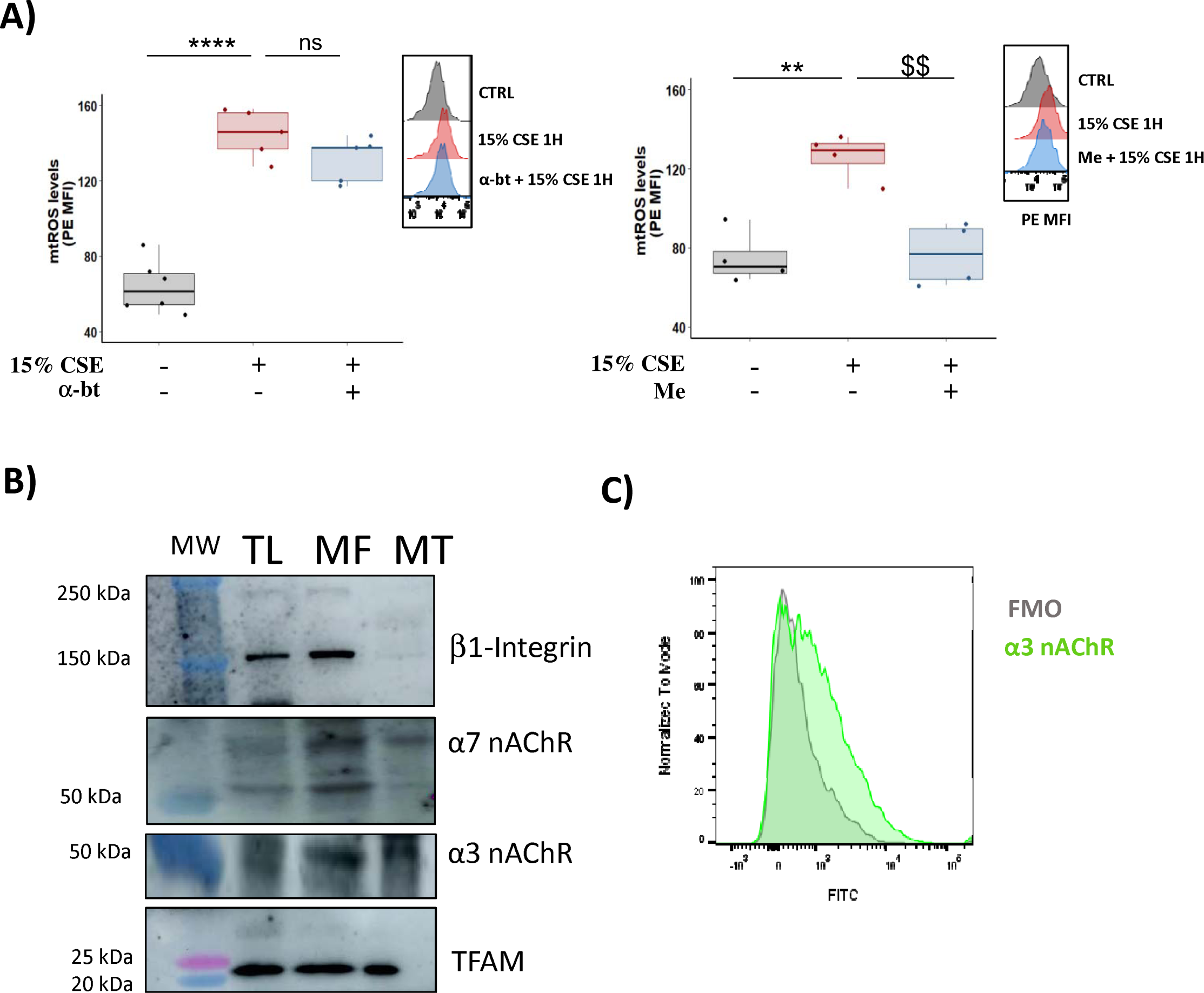
nAChR effects of nAChR antagonists on CSE-induced rise in mtROS and expression in mitochondria. **A)** Cells were pre-treated with 100 nM α-bungarotoxin (α-bt, *left*) or 10 µM mecamylamine (Me, *right*) and then exposed to 15% CSE for 1 hour. After this time, mtROS levels were measured with mitoSOX probe and analyzed by flow cytometry. Data are shown as interquartile range (p75 upper edge of box, p25 lower edge, p50 midline) as well as the p95 (line above box) and p5 (line below); (n= 3-6). Representative half-overlay histograms of mitoSOX fluorescence intensity are shown: control non-CSE treated (CTRL, *grey*), 15% CSE (*red*), α-bt + 15% CSE (*blue*), me + 15% CSE (*blue*). Statistical comparison between groups was performed using one-way ANOVA test followed by Dunnet’s post hoc test; (**)p<0.01, (****)p<0.0001, (ns) non-significant. **B)** Western blot analysis of nAChR in isolated mitochondria from hPASMC. Representative images of total cell lysate (TL), membrane fraction (MF) and isolated mitochondria (MT) probed with anti-β1 integrin (to detect plasma membrane contamination), anti-α3 nAChR, anti-α7 nAChR, and TFAM (as a mitochondrial control marker); (n=3). **C)** α3-nAChR expression in mitochondria shown as FITC intensity; FMO= fluorescence minus one.

### Inhibition of nAChR prevents functional alterations and decreased PA reactivity mediated by CSE exposure

Our previously published results demonstrate that cigarette smoke directly contributes to pulmonary arterial remodeling through increased cell senescence (6). Therefore, we tested the ability of these nAChRs antagonists to reverse these effects, as well as those on cell cytoskeleton. The results showed that α-bt and mecamylamine were able to prevent CSE-mediated cellular senescence and mecamylamine prevented as well the increase on cell hypertrophy (Figure 7A). Additionally, the loss in actin cytoskeleton dynamics after CSE treatment was significantly prevented in the presence of these antagonists (Figure 7B), thus strengthening the link between channel activation, ROS production and cytoskeletal dysfunction.

**Figure 7.**
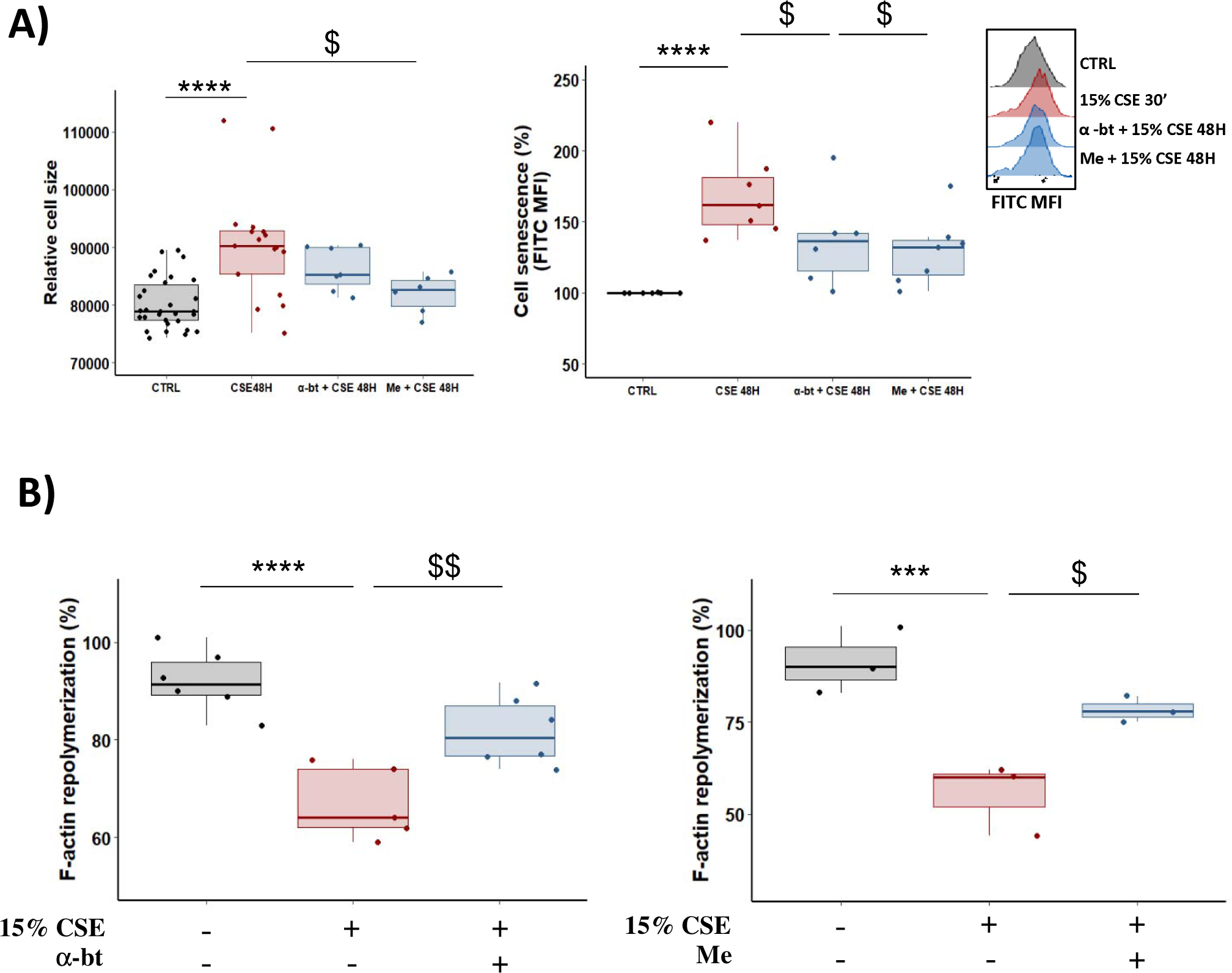
Effects of nAChR antagonists on functional outcomes induced by CSE exposure. **A)** Relative cell size and X-gal staining to quantify β-galactosidase activity were analyzed by flow cytometry. hPASMC were pre-treated with 100 nM α-bungarotoxin or 10 µM mecamylamine and then exposed to 15% CSE for 48 hours. Data on relative cell size were measured by forward light scattering and presented as interquartile range (p75 upper edge of box, p25 lower edge, p50 midline) as well as the p95 (line above box) and p5 (line below); (n_=_6-16). Cell senescence is presented as median fold change in FITC MFI and normalized to the levels in non-CSE treated (CTRL); (n=6-7). Representative half-overlay histograms of the median fluorescence intensity (MFI) are shown in control (*grey*), 15% CSE (*red*), α-bungarotoxin + 15% CSE (*blue*) and mecamylamine + 15% CSE (blue). **B)** Quantification of F-actin repolymerization percentage measured by flow cytometry. The repolymerization analysis was assessed in cells treated with 5 µM cytochalasin B for 2 hours and then allowed to recover for 1 hour in cells exposed to 15% CSE in the absence (-) or presence (+) of 100 nM α-bungarotoxin (*left*) or 10 µM Mecamylamine (*right*). The MFI of recovering cells was adjusted by the ratio between destabilized/non-destabilized cells. Data are shown as interquartile range (p75 upper edge of box, p25 lower edge, p50 midline) as well as the p95 (line above box) and p5 (line below) of the cell senescence percentage; (n= 3-6). Statistical comparison between groups was performed using one-way ANOVA test followed by Dunnet’s post hoc test. ns= not significant, (*)p<0.05, (**)p<0.01, (***)p<0.005, (****)p<0.0001 points to significance between control group and CSE-exposed group. ($)p<0.05, ($$)p<0.01 points to significance between CSE-exposed group and the respective antagonist treatment + CSE-exposed group.

To evaluate the functional contribution of nAChRs to vascular functions in a more physiological approach, we performed wire myography experiments with isolated pulmonary arteries from mice and treated them with CSE for 12h in the presence or absence of α-bt or mecamylamine. Our results proved that these inhibitors effectively halted the decline in endothelium-dependent vasodilation prompted by Ach or that induced by riociguat (Figure 8). These results underscore the importance of these channels in pulmonary artery vascular responses to tobacco smoke.

**Figure 8.**
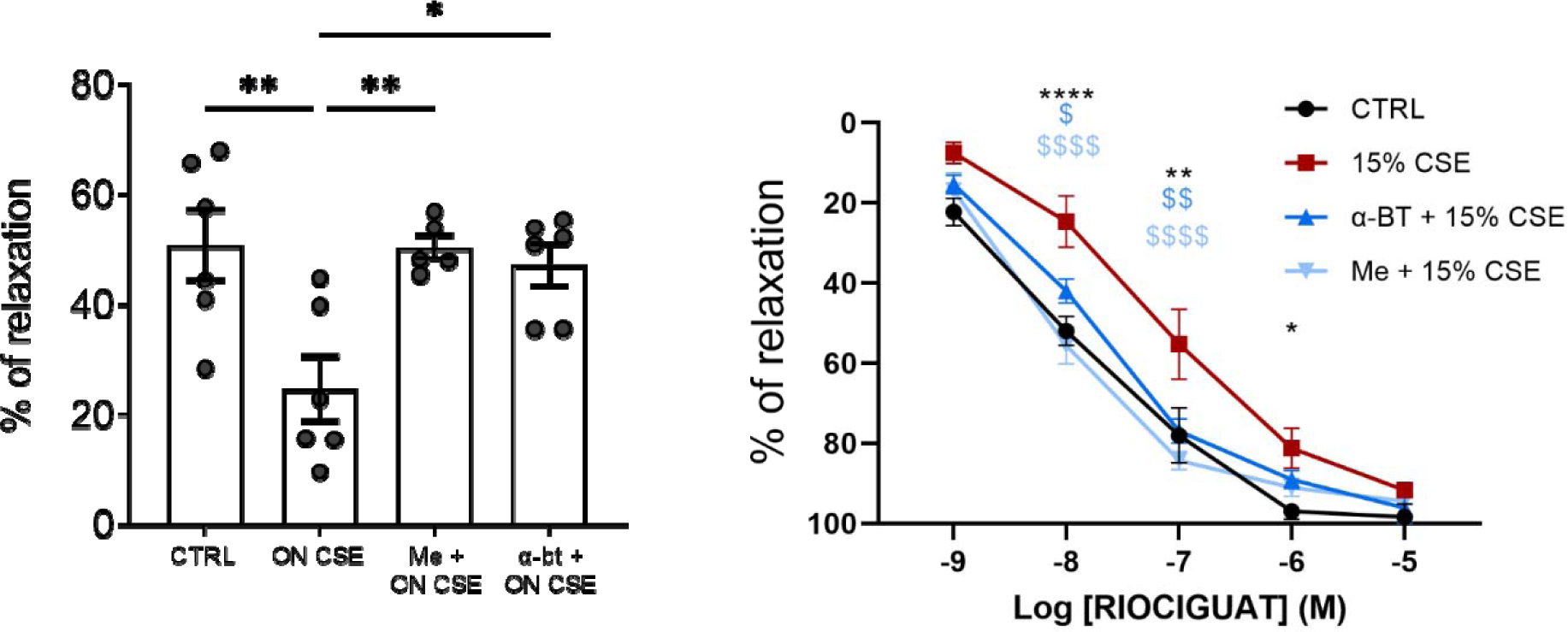
Effects of nAChR antagonists on PA reactivity ex vivo. Vascular responses were analyzed in endothelium-intact PAs from C57BL/6J mice (n=4/each group) previously incubated overnight with 15% CSE. Average values of the endothelium-dependent vasodilation induced by 10^-5^ M ACh (*left*) or increasing amounts of riociguat (*right*) were assessed in non-CSE treated (CTRL) (n_= 5 rings) or 15% CSE (n_=_4 rings) alone or in combination with mecamylamine (Me+CSE) or α-bungarotoxin (α-bt+CSE) and expressed as percentages of relaxation related to the maximum 5-HT-driven vasoconstriction. Statistical comparison between groups was performed using one-way ANOVA test followed by Dunnet’s post hoc test. (*)p<0.05, (**)p<0.01, (****)p<0.0001 points to significance between control group and CSE-exposed group. ($)p<0.05, ($$)p<0.01, ($$$$)p<0.0001 points to significance between CSE-exposed group and the respective antagonist treatment + CSE-exposed group.

## DISCUSSION

COPD related PH is a significant health concern with few treatments and poor outcomes. Our previous studies identified direct effects of tobacco smoke on the pulmonary vascular dysfunction, mostly concerning to vasodilation responses (6, 10). However, the mechanisms involved in CSE-mediated decrease on pulmonary arterial contractility are poorly understood. Our current studies delve into the molecular pathways related to the generation of ROS and the deregulation of calcium levels in pulmonary artery smooth muscle cells after their exposure to tobacco smoke, and how these pathways explain the smoke-induced decrease in pulmonary artery contractility.

Our findings demonstrate that exposure to CSE promoted oxidative stress in hPASMC in parallel with an increase in intracellular calcium, as well as alterations in the organization of proteins that contribute to the contractile function of the cell. The expression of nicotinic receptors in the human pulmonary artery, along with the results from using inhibitors or interfering with their expression, reveals for the first time the involvement of these receptors in the effects of tobacco on the human pulmonary artery. In this respect, several studies suggest that nicotine (31), tobacco smoke exposure (32) and particularly the increase in ROS (33, 34), have significant impacts on cytoskeleton structure, focal adhesions, and the dynamics of actin and tubulin cytoskeleton. Our results proved that hPASMC exposed to CSE for 24 hours had a disorganized cytoskeleton with decreased levels of several cytoskeletal proteins, these including tubulin, cortactin and vinculin. We also observed alterations in actin dynamics and MLC levels of phosphorylation, important hallmarks in cell contractility alterations. These effects seemed to be closely associated with the CSE-induced rise in oxidative stress, as evidenced by the response to treatment with the antioxidant NAC. In this respect, previous studies proposes that MLCK could be influenced by the cellular redox state and heightened oxidation levels might inhibit its activity, consequently impeding MLC phosphorylation (35). This could explain the lower arterial reactivity observed in this study.

Previous studies have underscored the interplay between ROS and calcium, important intracellular signaling molecules governing various physiological and pathophysiological processes, these including vascular pathologies (36). Our results demonstrated an elevation of intracellular calcium levels that correlated with a subsequent increase in total and mitochondrial ROS in hPASMC after CSE exposure. These findings hold significant importance, as they suggest that calcium serves as the primary signaling element in CSE-mediated effects. Furthermore, it indicates that the initial surge in cytoplasmic calcium levels directs the dysregulation of redox balance. Furthermore, it underscores the intricate interplay between ROS and calcium and emphasizes their involvement in vascular contractility dysfunction following short-term CSE exposure. In this context, Song and colleagues described a positive reciprocal loop connecting these two elements in the context of pulmonary hypertension (37). This model supports the main role of calcium in ROS production after CSE exposure. Despite there seems to be some interaction between calcium and mtROS, their exact relationship and influence on mitochondrial dysfunction are not yet fully understood. In this context, prior research emphasizes the interplay between mtROS and calcium elucidating how alterations in calcium channel activity modulates mitochondrial calcium which in turn promotes the increase on mtROS (38, 39). This stablished relationship between the calcium release by the ER and the production of mtROS is in line with our findings suggesting a potential interaction between mtROS and calcium in cells treated with CSE. Our results indicate that this signaling interaction has the potential to shift toward a pathological condition and thus would have implications for pulmonary arterial dysfunction secondary to COPD.

nAChR are widely distributed in neuronal and non-neuronal tissues, where they play diverse physiological roles. Recently α3, α4 and α7 nAChR have been identified in the bronchial and respiratory tract (40, 41, 42). These receptors, in particular α7nAChR, appear to have protective effects against chronic inflammation, thus offering therapeutic benefits in the setting of pulmonary emphysema (43, 25, 44). However, their effects may vary across different types of airway cells and under exposure to exogenous stimuli such as nicotine (45). In this respect, previous studies highlight the deleterious effects of these channels in airway remodeling and dysfunction (41, 24). Thus, inhibition of α7 nAChRs may also represent a novel target in patients with airway diseases. Despite cigarette smoking is an important risk factor for the development of vascular diseases, and nicotine induces vascular dysfunction (46), research on nAChR expression in pulmonary vascular system is still limited. Furthermore, the impact of nicotine exposure on the cardiovascular system is complex, as its direct effects differ according to cell type. Previous studies identified multiple nAChR alpha-subunits in rat arteries (47, 48) and there is strong evidence that nicotine exerts deleterious effects on the vasculature through its interaction with nAChRs (49), being α7 nAChRs a key player in these processes (50). These results are in line with previous studies in which α7 nAChR is the transducer receptor for nicotinic signaling. However, our present results in human pulmonary artery cells proved that both nAChR α3 and α7 are involved. Additionally, previous studies have shown that nicotine induces reorganization of cytoskeletal structures in vascular SMC, including changes in α-actin, vimentin, and β-tubulin, and vascular dysfunction. These have been shown to be at least partially mediated by the α7 nAChR (51, 52, 53). On the other hand, α7 nAChR exhibits high permeability to calcium, enough to couple the activity of this receptor to intracellular calcium signaling pathways (54). In line with this, our results using pure nicotine and the nAChR antagonists, α-bungarotoxin and mecamylamine, as well as the specific interference from the α3 or α7 nAChRs, indicated a direct role of both receptors in the accumulation of intracellular calcium and oxidative stress mediated by CSE in the pulmonary artery smooth muscle cells. Furthermore, our results also proved detectable levels of α3 and α7 nAChRs in the mitochondria, which may also contribute to CSE-mediated effects. These receptors have been previously found in the outer mitochondrial membrane, and nicotine-mediated activation depolarize it, promoting calcium discharge from endoplasmic reticulum (ER) via receptors located in this organelle, this leading to elevated oxidative stress within the mitochondria (55, 56, 38). This process highlights the reticulum-mitochondria axis as a pivotal component of the CSE-mediated effects on mitochondrial dysfunction in hPASMC.

While previous studies have suggested the potential of nAChRs agonists in COPD inflammation, there is no previous study confirming their expression in the human pulmonary artery. Our current research is the first to unveil the presence of nAChRs in human pulmonary artery and to elucidate their biological significance in CSE-induced impairment of pulmonary artery function. Activation of these receptors by nicotine triggers calcium signaling pathways and promotes oxidative stress. These events alter the contractile machinery and actin dynamics within human pulmonary artery SMCs, collectively contributing to pulmonary artery dysfunction. Most importantly, blocking α3 and α7 nAChR effectively prevented the decrease in vasodilation stimulated by Ach or riociguat in PA challenged with CSE. These findings hold significant clinical relevance, indicating that treatment with inhibitors targeting nicotinic acetylcholine receptors (nAChRs), alongside existing vasodilators like riociguat, could potentially provide a therapeutic alternative for patients with COPD-related pulmonary hypertension who do not respond to this medication.

